# Conformational Dynamics and metal-ion interactions in human small Ubiquitin-like modifier (SUMO2)

**DOI:** 10.1101/2023.12.19.572351

**Authors:** Anupreet Kaur, Harpreet Singh, Dinesh Kumar, Gagandeep Kaur Gahlay, Venus Singh Mithu

## Abstract

SUMO (Small Ubiquitin-like Modifiers) proteins are involved in a crucial post-translational modification commonly termed as SUMOylation. Currently little is known about localisation and accumulation of SUMO2 conjugates in response to proteasome inhibitors. In this work, we have investigated a strong binding of Cu^2+^ ions in the C-terminal region of SUMO2 resulting in its aggregation and seems to interefere in it’s non-covalent interaction with a V/I-X-V/I-V/I based SIM. In Ubiquitin-proteasome system, SUMO2 controls many targets in regulation of all aspects of metabolism. The conformational flexibility of SUMO2 could play a crucial role, therefore we have also characterized native-state flexibility of human SUMO2. We show that compared to SUMO1, several amino acids in SUMO2 around α_1_-helix region access energetically similar near-native conformations. This could play a fundamental role in its non-covalent interactions with SUMO interaction motifs (SIMs).

## 1. Introduction

Covalent tagging of proteins by SUMO proteins, also known as SUMOylation, is a crucial post-translational modification [1]. It is mechanistically similar to Ubiquitination and involves the covalent isopeptide bond linkage to Lysine residue in ψ-K-x-D/E motif in target molecules [2]. SUMOylation pathway and its consequences involve numerous covalent and non-covalent interactions. SUMOylation can alter protein function by blocking the old, or by exposing the new interaction sites [3]. Highly expressed isoforms of SUMO proteins in humans are identified as SUMO1, 2, and 3. The localization and function of these proteins within cells make SUMO paralogs dissimilar to each other. SUMO1 is found on the nuclear envelope and cytoplasm whereas, SUMO2/3 lies near to chromosomes [4]. SUMO1 and SUMO2 have both distinct and overlapping sets of substrates, indicating their role in redundant and non-redundant cellular functions [5-9]. In events like embryogenesis, SUMO2 is the predominantly expressed isoform and can compensate for the functions of SUMO1 and SUMO3 [10]. Despite having a similar Ubiquitin-like fold, SUMO2 shares only 45% sequence identity with SUMO1 [11]. Differences occur mostly in the β_2_-strand and the α–helix, the structural region which is known to bind non-covalently with the SUMO interaction motifs in target molecules [12].

In contrast to SUMO1, SUMO2/3 conjugation rapidly increases upon proteotoxic stress to resolve the misfolding and insoluble protein aggregates [13-15]. The early conjugation of SUMO2/3 might ensure the protein solubility to minimise the misfolded aggregation [16]. The SUMO2/3 modification and Ubiquitin –proteosomal system (UPS) are strongly integrated and act in a cooperative way[17] [18]. The direct association between SUMO2/3 and Ubiquitin is also examined under focal cerebral ischemia [19]. Therefore, SUMO2/3 binding greatly contributes in regulation of degradation of proteins and its dysregulation might be associated with UPS failure [20]. The dysfunction of UPS may result in accumulation of many proteins and involve in the origin of many neurodegenerative diseases such as Alzheimer disease (AD)[21].

The Copper and Zinc complexes are recently discovered new UPS inhibitors, which induce aggregation and decrease the thermal stability of ubiquitin [22] [23] and SUMO1[24]. The Cu^2+^ and Zn^2+^ imbalance play paramount role by affecting protein structure and oxidative stress[25]. There is evidence that Cu dyshomeostasis leads oxidative stress in AD patients resulting in memory deficits[26].Similarly, the chelation of Zn^2+^ may have critical role in synaptic and memory deficits[27]. However, the binding properties of SUMO2 is still uncharacterized in presence of these inhibitors.

In this paper, we have characterized the binding of Cu^2+^ and Zn^2+^ ions with SUMO2 using fluorescence spectroscopy, gel electrophoresis, and NMR spectroscopy. The impact of metal ion binding on its ability to interact non-covalently with a V/I-x-V/I-V/I based SIM from Daxx protein is also examined. The dynamic nature of SUMO2 might enable the protein to regulate cellular processes during stress and reversible SUMOylation process make it to act in transient manner. Here, In this work, we have also quantified the native-state conformational flexibility of SUMO2 using the non-linear temperature dependence of amide proton chemical shifts (^1^H_N_δ). The temperature profiles were fitted to a two-state exchange model to determine flexible regions in the native state of SUMO2.

## 2. Material and methods

### 2.1. SUMO2 Cloning and Expression

#### 2.1.1. Cloning of Human SUMO2 in pQE-80L

SUMO2 cDNA encoding amino acids 1-92, corresponding to the active protein, was PCR amplified from pDONR221-SUMO2 plasmid (purchased from DNASU plasmid repository, United States) using Phusion® High-Fidelity DNA Polymerase (NEB, USA) with forward primer 5’ GATGGATCCATGGCCGACGAAAAG 3*’* and reverse primer 5’ TTCAAGCTTTTAACCTCCCGTCTGCTG 3*’* as per the manufacturer’s recommendations. The PCR product was digested with BamHI and HindIII, and ligated into BamHI-HindIII digested pQE-80L using T4 DNA ligase (NEB, USA).

#### 2.1.2. SUMO2 Protein Expression and Purification

The recombinant SUMO2 protein was over-expressed in *E. coli* BL21 DE3 cells with 1mM final IPTG concentration for 8 hours at 27°C, and purified using Ni-NTA column chromatography (Thermo Fisher Scientific, USA). Briefly, the induced pellet was re-suspended in sodium PBS buffer (50 mM NaH_2_PO_4_, 50 mM Na_2_HPO_4,_ 300 mM NaCl) supplemented with 10 mM Imidazole. The cells were sonicated for 10 minutes (5s pulse On; 5s pulse Off; 40% amplitude) on ice followed by centrifugation at 6000 rpm for 40 minutes. The clear supernatant containing recombinant protein was bound to Ni-NTA beads at pH 8 at 4*°*C. The beads were washed three times with 3 column volumes of wash buffer (sodium PBS buffer, 30 mM Imidazole). The recombinant protein was eluted four times with one column volumes of elution buffer (sodium PBS buffer, 300 mM Imidazole). For NMR studies, labeled SUMO2 protein was purified as above except that the recombinant cells were grown in M9 minimal media containing ^15^NH_4_Cl, and finally reconstituted in a buffer having 25 mM sodium phosphate (pH 5.5) containing 150 mM NaCl, 5 mM EDTA, and 1 mM DTT.

### 2.2. SIM Synthesis

The *Daxx12-H* (DPEEIIVLSDSD) was chemically synthesized in a solid phase peptide synthesizer (PS3; Protein Technologies, Medford, MA) as described previously[28].

### 2.3. Fluorescence spectroscopy

25 μM SUMO2 protein solution was used for fluorescence measurements in sodium phosphate buffer. PerkinElmer LS-55 spectrometer was used for all measurements using an excitation wavelength at 280nm (slit width=15nm) and the emission profiles recording between 290 and 600nm (slit width=7nm, scan speed=600nm/min) at 27*°*C. The metal stock solution (Zn^2+^ / Cu^2+^; 1M) was also prepared in phosphate buffer for protein-metal ion titrations. The required concentration of metal was added to a protein solution, followed by equilibration for 15 minutes. Fluorescence intensity (*F*) was monitored at 345nm as a function of metal ion concentration [*M*^2+^],. The binding constant (*Kb*)and number of binding sites (*n*) [29] were calculated using the modified Stern–Volmer equation as follows

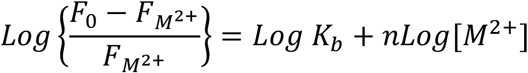

### 2.4. SUMO2 aggregation

14% Tris-Tricine non-reducing silver-stained gel was used to monitor concentration and time-dependent aggregation. 50μM SUMO2 protein was incubated for 30 minutes in the absence and presence of Zn^2+^ / Cu^2+^ ions (0.2-4.0 mole equivalents) at 27°C. Non-reducing buffer (12% Sodium lauryl sulfate, 30% glycerol, 0.05% Coomassie Brilliant Blue blue G-250, 150mM Tris/HCl) was added to the sample and heated at 95°C for 2 min to stop the reaction. The gel was stained with 1% silver nitrate to visualize different configurations of the protein [30]. For longer incubation studies, the sample containing an equimolar concentration of protein and metal ions was incubated at 27°C for 48 hours and an aliquot was drawn at different time intervals for electrophoresis and Bradford assay.

### 2.5. NMR Studies

#### 2.5.1. Temperature Dependence of Amide protons

NMR measurements were made using a 5 mm NMR tube (Wilmad Glass, USA) with a sealed capillary tube containing 0.1 mM DSS dissolved in deuterium oxide (D_2_O) for the locking and chemical shift referencing. All ^15^N-^1^H heteronuclear single quantum correlation (HSQC) spectra were recorded on an 800MHz Bruker spectrometer using 300 time points in the indirect dimension with a dwell time of 39 μs. All NMR data were processed using Topspin3.6.0 (Bruker) and analyzed using Analysis CCPNMR (2.3.1) [31]. **Fig. S1** shows the ^15^N-^1^H HSQC spectrum of SUMO2 along with the peak assignments as assigned by BMRB format (entry number-25577).

#### 2.5.2. Backbone Dynamics

^15^N relaxation measurements (^15^N*-*T_1_, ^15^N*-*T_2_, and {^1^H}*-*^15^N*-*NOE) were performed at 11.74 T and 18.79 T corresponding to _1_H Larmor frequency of 500 and 800 MHz, respectively. A set of ^1^H-^15^N HSQC spectra were recorded for T_1_ (10, 40, 80, 160, 640, 1240, 1920 ms), T_2_ (20, 60, 100, 140, 180, 220, 260 ms) measurements at 500 MHz, and T_1_ (10, 50, 70, 110, 170, 270, 350, 550, 770, 990 ms), T_2_ (18.56, 37.12, 55.68, 74.24, 92.8, 111.36, 129.92 ms) measurements at 800 MHz. Intensity of 82 unambiguous peaks was fitted to the equation I_t_ = I_0_e^−t/T^ where I_*t*_ corresponds to an intensity at a given time point t, and T corresponds to extracted relaxation times (T_1_ and T_2_). The spectra are recorded in the presence and absence of ^1^H saturation for NOE measurements. Combined T_1_, T_2,_ and NOE data from two spectrometers were analyzed based on a model*-*free approach using Dynamic center (Bruker BioSpin GmbH).

#### 2.5.3. Protein-SIM interaction

100 mM stock solution of Daxx12 peptide (SIM) was prepared in DMSO. 500 μM ^15^N labeled SUMO2 protein was titrated using 0.2-2.0 mole equivalents of SIM while recording ^1^H-^15^N HSQC spectrum at each titration point. The chemical shift perturbation (CSP) between the bound and free form was calculated using the following equation.

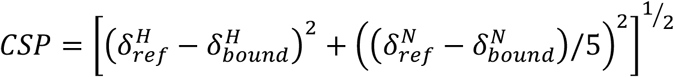

Where *δ*^*H*^and *δ*^*N*^ are the chemical shifts of amide proton and nitrogen, respectively. CSPs observed for control samples containing only protein and DMSO were subtracted from each titration point.

#### 2.5.4. Protein-Cu^2+^ ion interaction

The ^1^H-^15^N HSQC spectrum of 500 μM SUMO2 solution containing Cu^2+^ ions in desired concentration (0.0, 0.2, 0.4, 0.6, 0.8, 1.0 mole equivalents) was recorded by collecting 128 points in indirect dimension using a dwell time of 71.4 μs. Each spectrum was processed using Topspin3.6.0 (Bruker) and analyzed using Analysis CCPNMR (2.3.1) [31].

## 3. Results

### 3.1. SUMO2-Metal ion Interaction

#### 3.1.1. Binding affinity and stability of SUMO2

The metal ion binding affinity of SUMO2 was determined by monitoring the quenching of its fluorescence emission maxima (*λ*_max_=345 nm) upon the addition of Cu^2+^ and Zn^2+^ ions. **Fig. 1a** shows the plot of 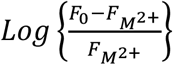 versus *log[Q]* for Cu^2+^ ions as per Stern-Volmer model used to calculate binding constant (*K*_*a*_ = 1.16 *×* 10^6^ M^-1^). As compared to Cu^2+^, no significant quenching was observed in the case of Zn^2+^ ions indicating a lack of interaction with SUMO2.

**Figure 1.**
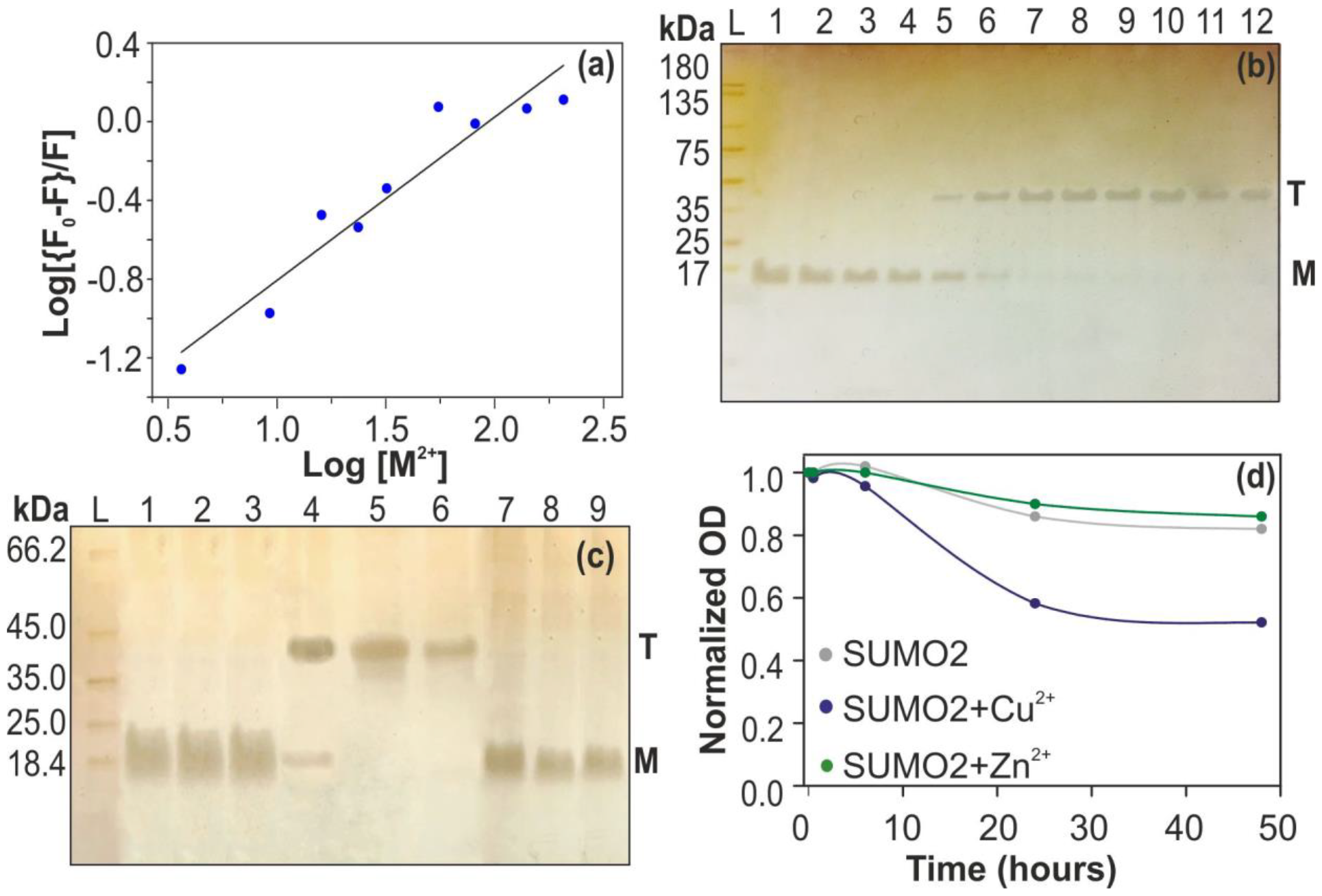
Modified stern-Volmer plot showing quenching of SUMO2 fluorescence as a function of Cu^2+^ concentration. Solide line represents linear fitting used to obtain binding constant (K_a_) and binding sites (n_b_). (b) Silver stained 14% non-Reducing Tricine SDS-PAGE showing concentration dependent impact of Cu^2+^ on SUMO2. SUMO2 was incubated for 30 minutes in the presence of 0.00, 0.20, 0.40, 0.60, 0.80, 570 1.00, 1.25, 1.50, 2.00, 2.50, 3.00, and 4.00 mole equivalents of Cu^2+^ (Lanes 1 to 13, respectively). (c) Impact of Cu^2+^ and Zn^2+^ ions on SUMO2 as a function of time. SUMO2 alone (Lane 1-3) incubated for 30 min (L1); 24h (L2); 48h (L3). SUMO2 incubated with an equimolar concentration of Cu^2+^ (Lanes 4-6) and Zn^2+^ (Lanes 7-9) for 30min (Lane 4 and 7), 24h (Lane 5 and 8), and 48h (Lane 6 and 9). Lane L corresponds to the unstained protein marker. Letters M and T are used to highlight the monomeric and trimeric states of SUMO2, respectively. (d) Normalized optical density OD_595_ as a function of incubation time measured using the Bradford assay.

**Fig. 1b** shows SDS-PAGE of 1.0-mole equivalent SUMO2 when incubated (for 30 min) with increasing Cu^2+^ ion concentrations. SUMO2 shows a single band corresponding to its monomeric state (∼12 kDa) up to 0.6-mole equivalents of Cu^2+^ (Lanes 2-4). At 0.8 mole-equivalents, an additional band corresponding to a trimeric state (∼35 kDa) appeared in the SDS-PAGE (Lane 5). This band continues to grow stronger as more Cu^2+^ is added (Lane 6), while the monomer band disappears at and after the addition of 1.5-mole equivalents of Cu^2+^ (Lanes 7-12). We also monitored the effect of longer incubation periods on the aggregation propensity of SUMO2 in presence of an equimolar amount of Cu^2+^ ions. **Fig. 1c** shows non-reducing SDS-PAGE of SUMO2 in the absence (Lanes 1-3) and presence (Lanes 4-6) of Cu^2+^ ions and incubated for 30 min (Lanes 1 and 4), 24 hr (Lanes 2 and 5), and 48 hr (Lanes 3 and 6). In presence of Cu^2+^ and at longer incubation periods, the amount of trimer gets reduced and large SUMO2 aggregates which are unable to enter the separating gel get formed. Standard Bradford assay was used to quantify the amount of these non-soluble aggregates of SUMO2 (**Fig. 1d**). It is found that nearly 50% of SUMO2 exists in form of non-soluble aggregates when incubated with an equimolar amount of Cu^2+^ ions for 48 hrs at 27*°*C. In contrast, SUMO2 remains largely monomeric under the influence of Zn^2+^ ions up to 4.0-mole equivalents, and at longer incubation periods with 1.0-mole equivalents of Zn^2+^ (Lanes 7-9, **Fig. 1c**). Also, it doesn’t show any aggregation enhancement in form of non-soluble aggregates (**Fig. 1d**).

#### 3.1.2. Cu^2+^ binding sites in SUMO2

We next looked into the binding site of Cu^2+^ using ^15^N isotopically enriched SUMO2. ^15^N-^1^H cross-peaks in the HSQC spectrum of SUMO2 recorded at 300 K were assigned using chemical shift values provided in BMRB assignment (ID= 25577). **Fig. 2a** shows an overlay of ^15^N-^1^H HSQC spectra of SUMO2 recorded in the absence and presence of 1.0-mole equivalents of Cu^2+^ at 300K (pH=6). Being paramagnetic, Cu^2+^ quenches NMR signals of spatially proximal nuclei, a property commonly used to identify Cu^2+^ binding sites in proteins. The relative signal intensity of each amide proton (I) compared to that observed in absence of Cu^2+^ ions (I_0_) was calculated and is plotted in **Fig. 2b**. Nearly each amino acid experiences quenching (average I/I_0_ = 0.57±0.31) in SUMO2 due to specific or non-specific interaction with Cu^2+^ ions. However, two specific sites show maximum signal quenching (I/I_0_ < av-σ). The first one is in the N-terminal and involves amino acids M1, D3, E4, and K5. The other site is located in the loop region preceding the β_5_-sheet leading to significant signal quenching of amino acids E77, M78, E79, D80, E81, D82, and T83. A closer look into the structural arrangement of these amino acids (**Fig. 2c**) shows that spatial proximal side chains of E79, D80, E81, and D82 could be involved in Cu^2+^ complexation. Three other amino acids; K35, D63, and Q89 also show significant quenching of NMR signals. However, a lack of similar amino acids in their close vicinity, sequential or spatial, tells us that that the observed quenching might be resulting from non-specific interactions.

**Figure 2.**
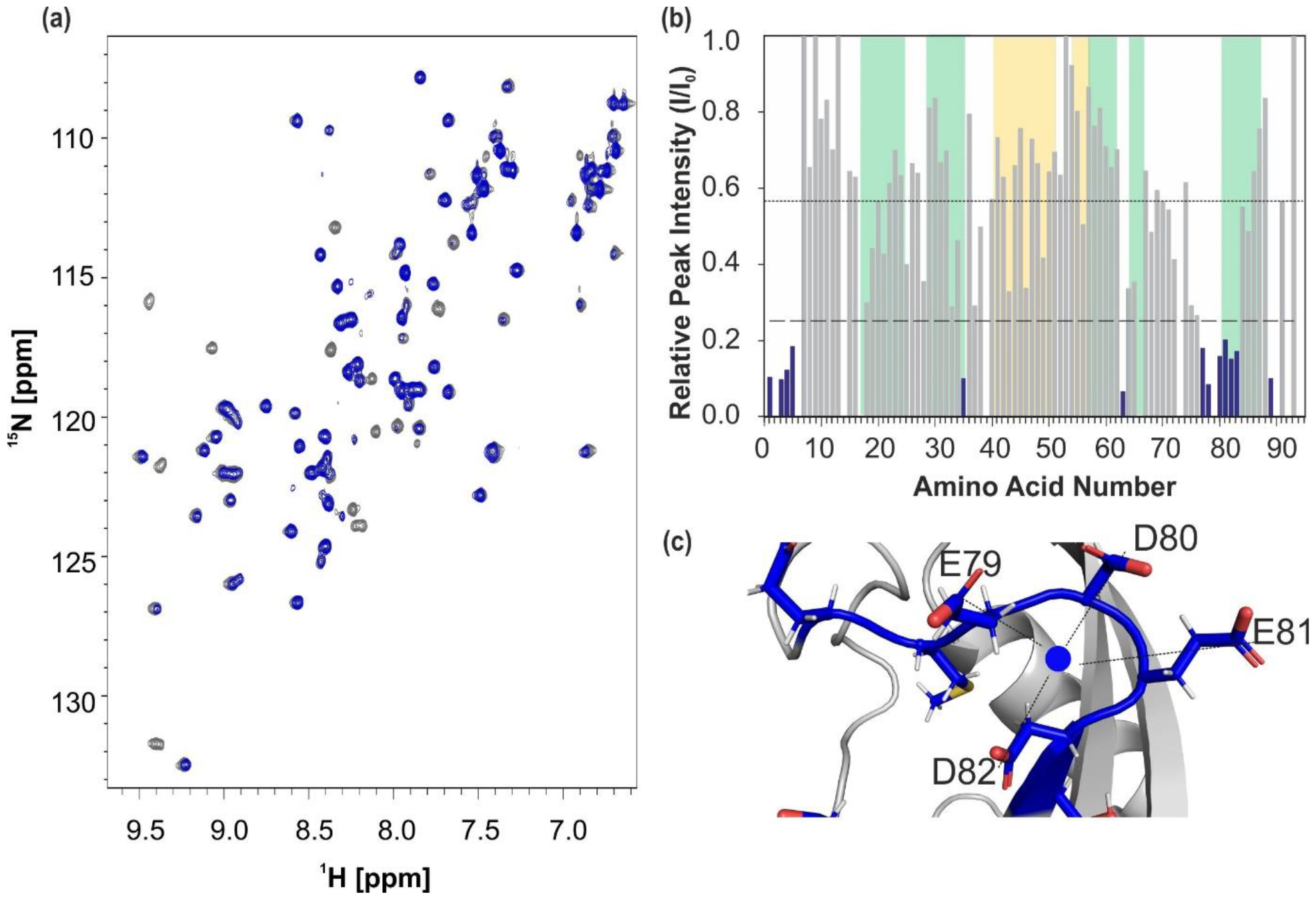
(a) An overlay of ^15^N-^1^H HSQC spectra of SUMO2 recorded in the absence (grey peaks) and presence of 1.0-mole equivalents of Cu^2+^ (blue peaks) at 300K (pH=6). (b) Relative intensities of ^15^N-^1^H cross-peaks in HSQC spectra of SUMO2 in 0 presence of 1.0-mole equivalents of Cu^2+^. Average quenching 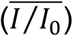 and standard deviation (*σ*) are shown using short- and long-dashed horizontal lines, respectively. Amino acids experiencing significant signal quenching 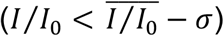 are highlighted with blue-filled bars. The β-sheet and α-helix structural regions of the protein are represented by green and yellow coloured regions. (c) The putative interaction site of Cu^2+^ with side chains of E79, D80, E81, and D82.

### 3.2. Impact of Cu^2+^ binding on SUMO2-SIM interactions

As such, Cu^2+^ binding doesn’t cause any significant change in the chemical shifts of amino acids in SUMO2. This indicates that the secondary structure of SUMO2 remains preserved in the event of its interaction with Cu^2+^. However, one cannot rule out subtle but significant changes occurring in the tertiary fold and dynamics of SUMO2 which might have an impact on its activity. Moreover, Cu^2+^ induced self-association of SUMO2 could also interfere in its interaction with the target molecules. To verify this, we have studied its interaction with Daxx12 peptide containing a [V/I]-X-[V/I]-[V/I] based SIM from human Daxx protein (^732^E**IIVL**SDSD^740^) [32]. To characterize its interaction with Daxx12, a set of ^15^N-^1^H HSQC spectra was recorded using 0.2, 0.4, 0.6, 0.8, 1.0, and 1.2 mole equivalents of Daxx12. Systematic changes were observed in chemical shifts of some amino acids indicating changes in their secondary structure due to SUMO2-SIM interaction. To quantify these changes, compounded chemical shift perturbations (CSP) were calculated for each amino acid at an equimolar protein to peptide ratio and are shown in **Fig. 3a**. Based on these values, amino acids showing CSP were classified as weakly 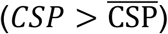 and strongly 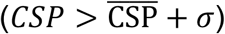 perturbed. These amino acids are highlighted on protein structure in **Fig. 3b**. As expected, β_2_-loop-α_1_ structural region in SUMO2 containing many such amino acids could be the possible interaction site of the Daxx12 peptide. This observation once again confirms that β_2_-loop-α_1_ structural region is always the choice of binding for such [V/I]-X-[V/I]-[V/I] based SIM. Since Cu^2+^ has a binding site on the opposite face of SUMO2, it is not expected to interfere directly in the SUMO-SIM interaction. However, minimal perturbations were observed when Daxx12 is added to SUMO2 solution containing an equimolar amount of Cu^2+^ ions (**Fig. 3a**). It seems the SUMO-SIM interaction is no longer happening when SUMO2 exists in a Cu^2+^ bound state.

**Figure 3.**
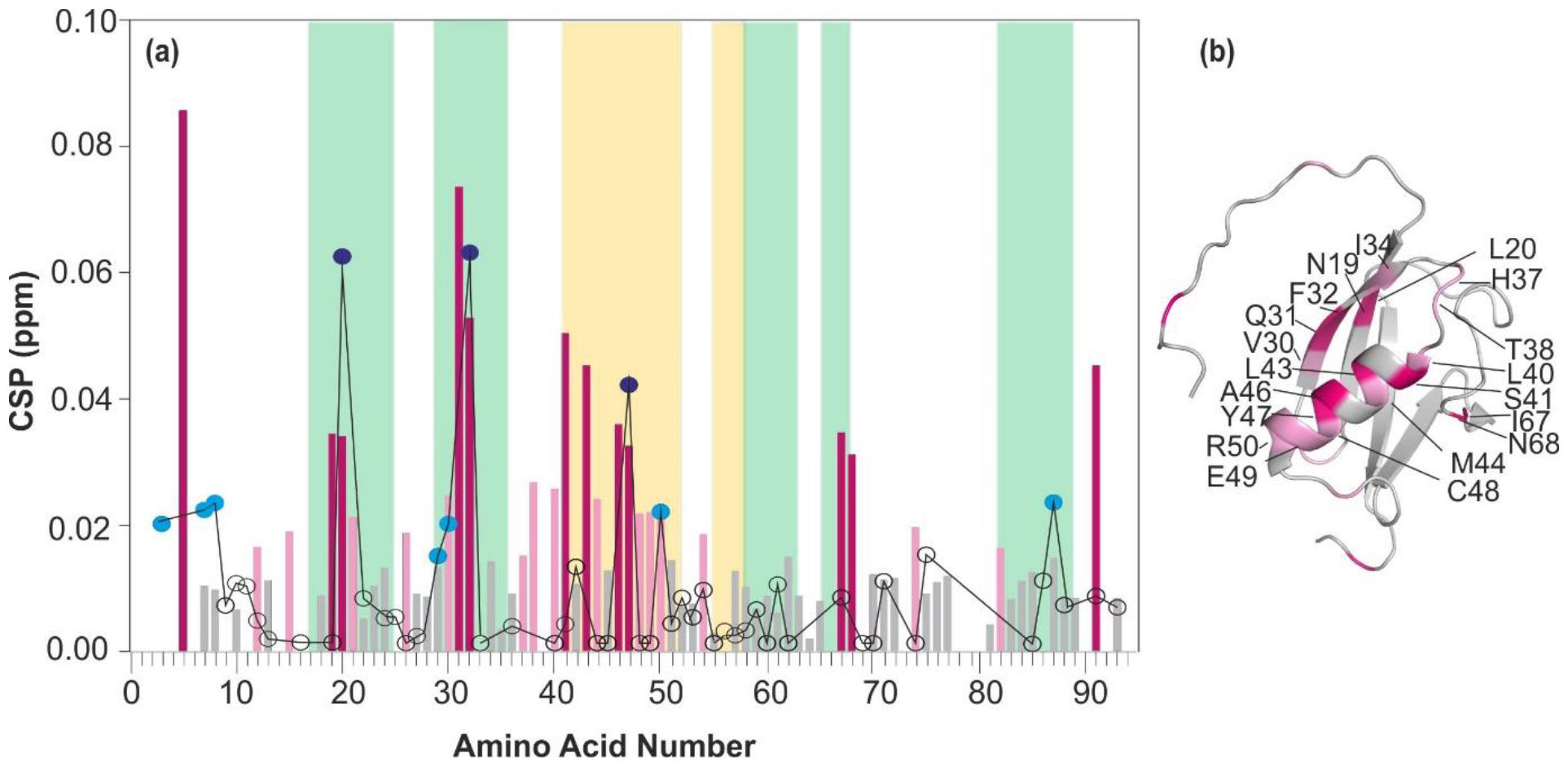
(a) Chemical shift perturbation observed upon interaction with Daxx12 for each assignable amino acid of SUMO2 in the absence (bars; grey (<av), light-pink (>av, <av+ σ), and deep-pink (>av+σ)) and presence (circles; hollow (<av), cyan filled circles (>av, <av+σ), and blue filled circles (>av+σ)) of Cu^2+^ plotted as a function of amino acid number. (b) The interacting residues and region of SUMO2 with Daxx12 are highlighted in the structural model.

### 3.3. Conformational fluctuations in the native state of SUMO2

The chemical shift of an amide proton is very sensitive to its hydrogen bonding network and varies according to changes in the secondary/tertiary structure of a protein molecule [33]. In general, it decreases linearly with a temperature rise [34] indicating elongation of the H-bond. The temperature coefficient varies with the structural location and can be used to differentiate between inter and intramolecular H-bonds. In some cases, amino acids which are part of local structural fluctuations yield non-linear temperature-dependent profiles [35]. First shown by Williamson and co-workers [36], this method has been applied to several proteins to characterize their near-native energy landscape [37, 38]. To identify such amino acids in SUMO2, we have recorded a set of 8 ^15^N-^1^H HSQC spectra at 288, 292, 296, 300, 304, 308, 312, and 316 K. The spectra were assigned following a systematic shift in each cross-peak. A well-dispersed HSQC spectrum obtained at each temperature indicates that SUMO2 does not undergo any global unfolding in the studied temperature range. The chemical shift of each amide proton (^1^H_N_*δ*) was plotted as a function of temperature and fitted to a straight line to calculate the fitting residuals. The residuals so obtained were evaluated for curvatures by fitting them to a parabolic equation (*Δ* ^*1*^*HNδ* =*a* + *bT*+ cT^2^) and using the value of quadratic coefficient (c) as a measure of curvature. Statistically significant values (p < .005) were used to calculate the average 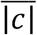 value (0.027 ± 0.030 ppm K^-2^), which was further used to identify weakly 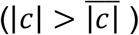 or strongly 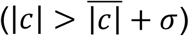 curved amino acids. Following this criterion, a total of 15 amino acids were identified as “curved” (6 weakly curved: Q13, G24, F60, D63, I67, and Q75, and 9 strong curved: N19, T38, L40, S41, L43, A46, C48, R61, and F62). Fitting residuals for these amino acids are shown in **Fig. 4**. Values of quadratic coefficient obtained for individual amino acid are shown in **Fig 5a**.

**Figure 4.**
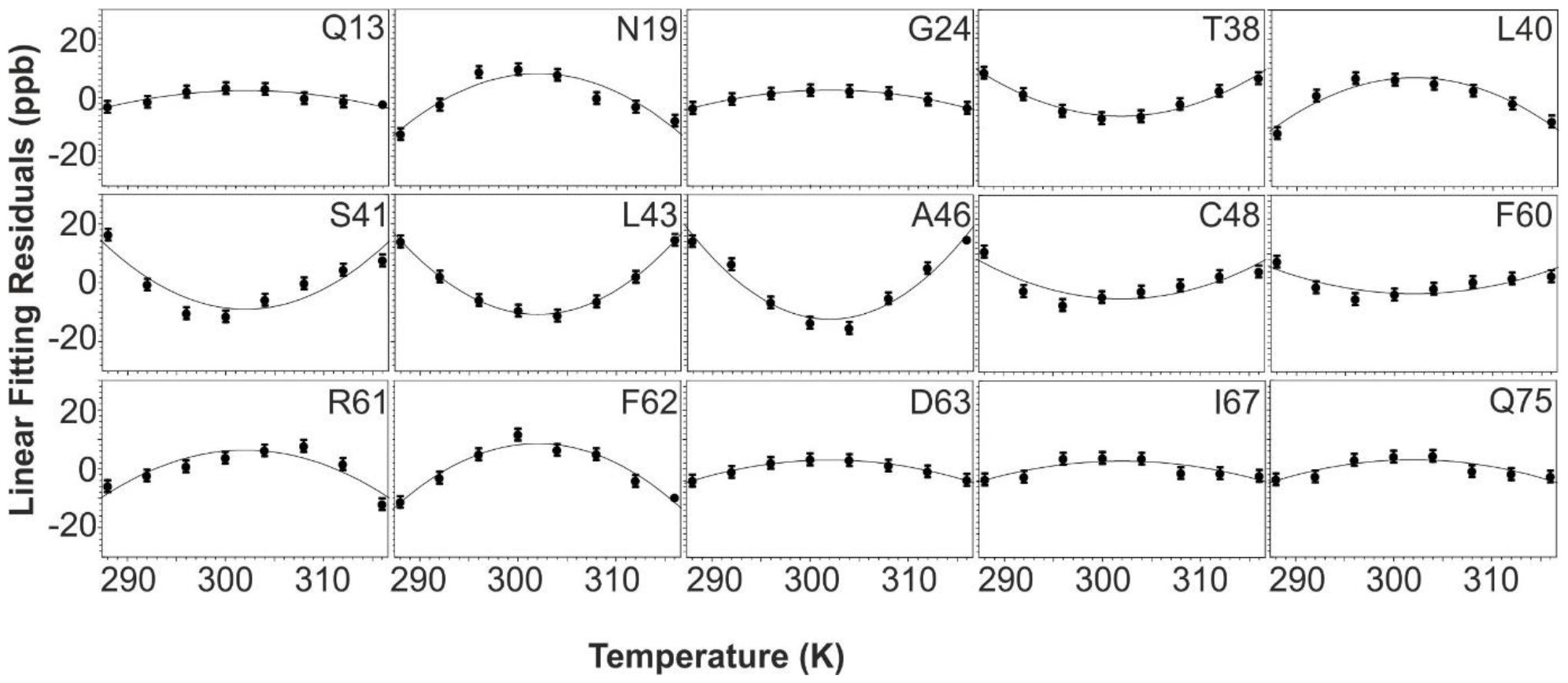
Fitting residuals for selective amino acids exhibiting a non-linear temperature dependence. The residuals are plotted against temperature and fitted to the parabolic equation (solid line) as described in the main text.

**Figure 5.**
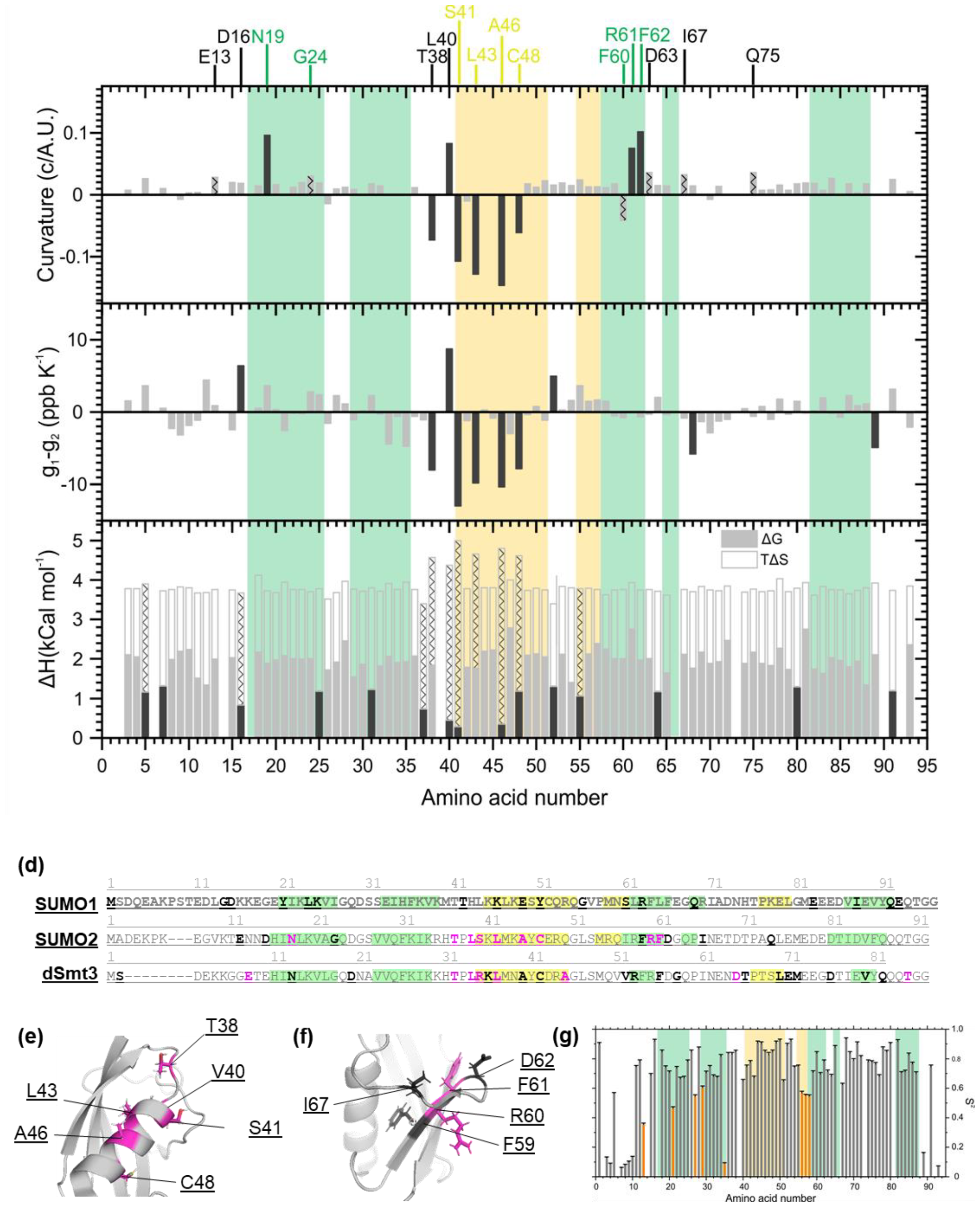
(a) Quadratic coefficient for assignable amino acids in SUMO2 as obtained from the quadratic fitting. (b) The difference in temperature coefficient of the ground (g_1_) and the near-native (g_2_) state as obtained from the two-state model fitting to temperature-dependent ^1^H_N_*δ*. (c) Other thermodynamic parameters as obtained from two-state model fitting. (d) Amino acids exhibiting strongly convex (in black) and concave (in pink) temperature profiles are highlighted on the sequence of SUMO1, SUMO2, and dSmt3. (e and f) Conformationally flexible amino acids in SUMO2 are highlighted on its structural model. (g) Square of the order parameter (S^2^) calculated using Lipari-Szabo model-free analysis of ^15^N-relaxation data (^15^N*-*T_1_, ^15^N*-*T_2_, and {^1^H}*-* ^15^N*-*NOE) of SUMO2. (a-d, g) β-sheet and α-helix structural regions of SUMO2 are highlighted using green and yellow colours, respectively.

The shape of curvature, either concave or convex, depends on the relative chemical environment of amide proton in the accessible near-native states i.e. their chemical shifts and temperature coefficients. On the other hand, the extent of curvature is dictated by the magnitude of difference in the thermodynamic parameters i.e. free-energy (ΔG), enthalpy (ΔH), and entropy (ΔS) [36]. To estimate these parameters, we have fitted the temperature profiles to a simple two state exchange model which calculates the observed chemical shift as

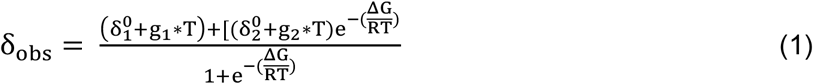

In this model, T is the temperature, δ^0^ is the chemical shift at absolute zero, *g* is the temperature coefficient, and *ΔG* represents the free-energy difference between the native (1) and near-native states (2) under fast-exchange. Changes incurred by chemical and thermodynamic parameters for each amino acid in SUMO2 are shown in **Fig. 5b** and **5c**, respectively. Out of the 15 curved amino acids, 6 (T38, S41, L43, A46, C48, and F60) exhibit concave curvatures. Except for F60, all of these show a significant negative change 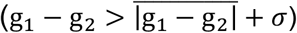 in temperature coefficient between the ground and alternative state. In general, the temperature coefficient of amide protons, which form an intra-molecular H-bond is more positive than those which make an H-bond with water [39]. Thus, amino acids with negative *g*_1_ − *g*_2_ values in **Fig. 5b** are the ones that are probably more exposed to water in their accessible near-native state. On the other hand, a strongly convex L40 incurs a positive change in temperature coefficient and thus seems to be more buried in the hydrophobic core in its near-native conformation. There are other amino acids (E13, N19, G24, F60, R61, F62, I67, and Q75) that are curved but do not show any significant change in temperature coefficient, and vice-versa (D16, G52, N68, and Q89). This could be because the temperature coefficient is mainly responsible for dictating the sign of curvature, but not its extent. Another reason for observed discrepancies could be the oversimplification of the near-native energy landscape using a two-state exchange model. Nevertheless, there seems to be a strong correlation between the sign of observed curvatures and the relative H-bonding environment of amino acids in the accessible near-native state.

Next, we analyzed the significant changes observed in the free energy 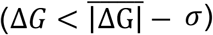, enthalpy 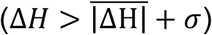, and entropy 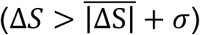 as shown in **Fig. 5c**. According to the model used here, an exchange between energetically similar (small Δ*G* value) and entropically/enthalpically dissimilar (large Δ*S* and Δ*H* values) near-native states results in strong curvatures [36]. Amino acids L40, S41, A46, and C48 satisfy all three conditions and accordingly exhibit strongly curved profiles. K5, D16, H37, T38, L43, and M55 meet on two conditions, and curved profiles were observed only in the case of T38 and L43 which incur significant change in enthalpy. While the occurrence of curvature ensures access to alternative conformations, the reverse is not always true. Nevertheless, it is quite clear that the α_1_-helix and the loop region connecting it to the β_2_-sheet are conformationally quite flexible. Six out of the nine strongly curved amino acids are located within a short stretch of eleven amino acids from T38 till C48 (**Fig. 5d & 5e**). This difference could have significant implications in the non-covalent interaction of SUMO proteins with V/I-x-V/I-V/I based interaction motifs which bind in the groove formed by β_2_-loop-α_1_ structural elements [40].

**Fig. 5f** highlights another conformationally flexible region in SUMO2, which is composed of amino acids F59, R60, F61, D62, and I67. Given the proximity of their bulky side chains, any local motion like ring flipping in the side chains of two phenylalanine residues could render this whole region conformationally flexible. Alternatively, it could be originating from an enhanced fast time-scale (ns-ps) backbone dynamics undergone by amino acids R56, Q57, and I57 in the close vicinity as indicated by their low order parameter values (**Fig. 5g**). Order parameters were calculated from ^15^N relaxation data (including T_1_, T_2,_ and {^1^H}-^15^N NOE) sets from two fields based on a model-free approach. Global isotropic rotational correlation time (*τ*_c_), calculated using T_1_/T_2_ ratios of selected residues, was found to be 7.02 ± 0.89 ns.

## 4. Discussion

Ubiquitination and SUMOylation are generally dependent protein modifications and extensive cross-talk exists between SUMO2 conjugation and Ubiquitin. The SUMO2 directly regulates the proteasome processing as well as indirectly involved in UPS system to provide sufficient free SUMO2 for further conjugation rounds[17]. Decrease in SUMO2 availability is sufficient to cause UPS dysfunction. Here, we show that Cu^2+^ ions target the two regions of SUMO2 and induce protein aagregation while no such binding is observed for Zn^2+^ ions.

In Comparison to SUMO1 isoform[24], Cu^2+^ ions share a strong interaction with SUMO1 (*K*_*a*_ = 1.31 *×* 10^6^ M^-^1) and promote its aggregation whereas Zn^2+^ has a weaker interaction with SUMO1 and doesn’t promote any aggregation. Cu^2+^ binds in the loop region preceding β_5_-sheet near the C-terminal. This region is rich in amino acids containing polar side chains (E83, E84, E85, and D86) capable of forming a coordination complex with Cu^2+^ ions. Cu^2+^ also binds SUMO2 at the same site which contains amino acids E79, D80, E81, and D82 at the equivalent locations (see section 3.2.2). Since this region is conformationally rigid in both SUMO1 and SUMO2 (**Fig. 5d**), this could be the reason why the binding site of Cu^2+^ remains conserved. Thus, Cu^2+^-SUMO interaction is conserved between these two human isoforms to a large extent. Zn^2+^ has a binding site in the loop-α_1_ region of SUMO1 involving amino acids M40, T41, T42, H43, and K46. Unlike SUMO1, this particular region in SUMO2 is rich with conformationally flexible amino acids and could be the possible reason behind the observed difference.

The Cu^2+^ and Zn^2+^ ion binding to SUMO proteins were demonstrated in our previous paper[24], However the exact effect of metal binding on the SIM based non-covalent interaction of SUMO is unexplained. To verify that Cu^2+^/Zn^2+^ interaction might affect its ability of SUMO to interact target proteins, we have performed NMR titration of Daxx12 peptide containing a [V/I]-X-[V/I]-[V/I] based SIM from human Daxx protein (^732^E**IIVL**SDSD^740^) with SUMO2 in presence and absence of Cu^2+^ ions. The results shows that β_2_-loop-α_1_ structural region is not sufficiently exposed for SIM interaction in the trimeric state of SUMO2 which is the predominant species under equimolar amount of Cu^2+^ ions. Additionally, Cu^2+^ binding could also result in allosteric structural changes in the β_2_-loop-α_1_ which can inhibit its binding with Daxx12.

Looking the native structure of SUMO2 for explaining its more stability over SUMO1 which is more aggregation prone in presence or absence of metal ions, SUMO2 shares a close similarity to dSmt3 (SUMO from Drosophilla melanogaster[41]) as both proteins were found to contain conformationally flexible amino acids at equivalent locations in the loop-α_1_ region (**Fig. 5d**). This includes amino acids T38, V40 S41, L43, A46, and C48 in SUMO2 (**Fig. 5e**) and T33, L35, R36, L38, A41, and C43 at equivalent locations in dSmt3. In SUMO2, most of these amino acids exhibit concave curvatures, an indication that they could be more water exposed in their alternative conformations. In other words, it is likely that in one of the near-native states accessed by SUMO2, the loop-α_1_ helix region is slightly opened up allowing for a relatively greater water interaction with these amino acids. The average order parameter value for structured amino acids (E13-Q88) in SUMO2 was found to be 0.75 ± 0.14 and is lower than that observed for SUMO1 indicating higher conformational disorder in SUMO2 due to fast time-scale motions [42]. E13, K21, G27, V29, K35, R56, Q57, and I57 are the only amino acids that yield significantly low order parameter values (<|*S*^2^| − *σ*). However, except for E13 which is weakly curved, none of the remaining were detected by the curved temperature dependence method as conformational flexible. There could be two reasons for this; (i) Near-native states accessed by these amino acids undergoing fast time-scale motions are chemically and/or thermodynamically quite similar, or (ii) Time-scale of motions undergone by amino acids with curved temperature profiles is not in ns-ps range.

## 5. Conclusions

We found that SUMO2-SIM interaction gets impaired in presence of Cu^2+^ ions. This could be because sub-stoichiometric amounts of Cu^2+^ are sufficient to drive SUMO2 into a trimeric state, which seems to evolve into higher aggregates with time. NMR studies indicate that Cu^2+^ binds in the C-terminal of SUMO2, possibly forming a coordination complex with amino acids E79, D80, E81, and D82 present in the loop region just before the β_5_ sheet. Conformational flexibility could play a crucial role in the functioning of a post-translation modifier like SUMO2. While sequence and structure render specificity to identify unique targets, conformational flexibility allows it to identify multiple targets. Our studies show that SUMO2 contains several amino acids in and around the α_1_-helix which adopt alternative conformations in the near-native landscape accessible to the protein. This conformational flexibility could play a crucial role in its non-covalent interactions with SUMO interaction motifs target molecules and is considered crucial for SUMOylation and its consequences. In a nutshell, our studies provide structural insights into the conformational flexibility in SUMO2 and its role in the interaction with [V/I]-X-[V/I]-[V/I] based SIMs and metal ions.

## CRediT authorship contribution statement

**Anupreet Kaur:** Conceptualization, Experimental work, Formal analysis, Writing - original draft. **Gourav:** Experimental work, Formal analysis. **Harpreet Singh:** Experimental work. **Gagandeep Kaur Gahlay:** Funding acquisition, Supervision, Writing - review & editing. **Venus Singh Mithu:** Conceptualization, Funding acquisition, Project administration, Methodology, Supervision, Writing - original draft, Writing - review & editing.

## Acknowledgements

VSM and GKG are thankful to the Department of Biotechnology, Ministry of Science & Technology, India for research funding (BT/ PR22289/BRB/10/1566/2016). Gourav is thankful to the Council of Scientific and Industrial Research, Ministry of Science & Technology, India for the junior research fellowship (file no. 09/254(0281)2018-EMR-I).

## References

1 Flotho, A. and Melchior, F. (2013) Sumoylation: a regulatory protein modification in health and disease. Annual review of biochemistry. 82

2 Gill, G. (2004) SUMO and ubiquitin in the nucleus: different functions, similar mechanisms? Genes & development. 18, 2046–2059

3 Kerscher, O. (2007) SUMO junction—what’s your function? EMBO reports. 8, 550–555

4 Ayaydin, F. and Dasso, M. (2004) Distinct in vivo dynamics of vertebrate SUMO paralogues. Molecular biology of the cell. 15, 5208–5218

5 Saitoh, H. and Hinchey, J. (2000) Functional heterogeneity of small ubiquitin-related protein modifiers SUMO-1 versus SUMO-2/3. Journal of Biological Chemistry. 275, 6252–6258

6 Azuma, Y., Arnaoutov, A. and Dasso, M. (2003) SUMO-2/3 regulates topoisomerase II in mitosis. The Journal of cell biology. 163, 477–487

7 Sternsdorf, T., Puccetti, E., Jensen, K., Hoelzer, D., Will, H., Ottmann, O. G. and Ruthardt, M. (1999) PIC-1/SUMO-1-modified PML-retinoic acid receptor α mediates arsenic trioxide-induced apoptosis in acute promyelocytic leukemia. Molecular and Cellular Biology. 19, 5170–5178

8 Kamitani, T., Nguyen, H. P., Kito, K., Fukuda-Kamitani, T. and Yeh, E. T. (1998) Covalent modification of PML by the sentrin family of ubiquitin-like proteins. Journal of Biological Chemistry. 273, 3117–3120

9 Mukherjee, S., Thomas, M., Dadgar, N., Lieberman, A. P. and Iñiguez-Lluhí, J. A. (2009) Small ubiquitin-like modifier (SUMO) modification of the androgen receptor attenuates polyglutamine-mediated aggregation. Journal of Biological Chemistry. 284, 21296–21306

10 Wang, L., Wansleeben, C., Zhao, S., Miao, P., Paschen, W. and Yang, W. (2014) SUMO2 is essential while SUMO3 is dispensable for mouse embryonic development. EMBO reports. 15, 878–885

11 Huang, W. C., Ko, T. P., Li, S. S. L. and Wang, A. H. J. (2004) Crystal structures of the human SUMO-2 protein at 1.6 Å and 1.2 Å resolution: implication on the functional differences of SUMO proteins. European journal of biochemistry. 271, 4114–4122

12 Song, J., Zhang, Z., Hu, W. and Chen, Y. (2005) Small Ubiquitin-like Modifier (SUMO) Recognition of a SUMO Binding Motif a reversal of the bound orientation. Journal of Biological Chemistry. 280, 40122–40129

13 Golebiowski, F., Matic, I., Tatham, M. H., Cole, C., Yin, Y., Nakamura, A., Cox, J., Barton, G. J., Mann, M. and Hay, R. T. (2009) System-wide changes to SUMO modifications in response to heat shock. Science signaling. 2, ra24–ra24

14 Hendriks, I. A., D’souza, R. C., Yang, B., Verlaan-de Vries, M., Mann, M. and Vertegaal, A. C. (2014) Uncovering global SUMOylation signaling networks in a site-specific manner. Nature structural & molecular biology. 21, 927–936

15 Tatham, M. H., Matic, I., Mann, M. and Hay, R. T. (2011) Comparative proteomic analysis identifies a role for SUMO in protein quality control. Science signaling. 4, rs4–rs4

16 Liebelt, F., Sebastian, R. M., Moore, C. L., Mulder, M. P., Ovaa, H., Shoulders, M. D. and Vertegaal, A. C. (2019) SUMOylation and the HSF1-regulated chaperone network converge to promote proteostasis in response to heat shock. Cell reports. 26, 236–249. e234

17 Schimmel, J., Larsen, K. M., Matic, I., van Hagen, M., Cox, J., Mann, M., Andersen, J. S. and Vertegaal, A. C. (2008) The ubiquitin-proteasome system is a key component of the SUMO-2/3 cycle. Molecular & Cellular Proteomics. 7, 2107–2122

18 Marinello, M., Werner, A., Giannone, M., Tahiri, K., Alves, S., Tesson, C., den Dunnen, W., Seeler, J.-S., Brice, A. and Sittler, A. (2019) SUMOylation by SUMO2 is implicated in the degradation of misfolded ataxin-7 via RNF4 in SCA7 models. Disease models & mechanisms. 12, dmm036145

19 Hochrainer, K., Jackman, K., Benakis, C., Anrather, J. and Iadecola, C. (2015) SUMO2/3 is associated with ubiquitinated protein aggregates in the mouse neocortex after middle cerebral artery occlusion. Journal of Cerebral Blood Flow & Metabolism. 35, 1–5

20 Liebelt, F. and Vertegaal, A. C. (2016) Ubiquitin-dependent and independent roles of SUMO in proteostasis. American Journal of Physiology-Cell Physiology. 311, C284–C296

21 Oddo, S. (2008) The ubiquitin-proteasome system in Alzheimer’s disease. Journal of cellular and molecular medicine. 12, 363–373

22 Arnesano, F., Scintilla, S., Calò, V., Bonfrate, E., Ingrosso, C., Losacco, M., Pellegrino, T., Rizzarelli, E. and Natile, G. (2009) Copper-triggered aggregation of ubiquitin. PLoS One. 4, e7052

23 Arena, G., Bellia, F., Frasca, G., Grasso, G., Lanza, V., Rizzarelli, E., Tabbi, G., Zito, V. and Milardi, D. (2013) Inorganic stressors of ubiquitin. Inorganic chemistry. 52, 9567–9573

24 Kaur, A., Jaiswal, N., Raj, R., Kumar, B., Kapur, S., Kumar, D., Gahlay, G. K. and Mithu, V. S. (2020) Characterization of Cu2+ and Zn2+ binding sites in SUMO1 and its impact on protein stability. International journal of biological macromolecules. 151, 204–211

25 Kozlowski, H., Luczkowski, M., Remelli, M. and Valensin, D. (2012) Copper, zinc and iron in neurodegenerative diseases (Alzheimer’s, Parkinson’s and prion diseases). Coordination Chemistry Reviews. 256, 2129–2141

26 Rossi, L., Squitti, R., Calabrese, L., Rotilio, G. and Rossini, P. (2007) Alteration of peripheral markers of copper homeostasis in Alzheimer’s disease patients: implications in aetiology and therapy. The journal of nutrition, health & aging. 11, 408

27 Watt, N. T., Whitehouse, I. J. and Hooper, N. M. (2011) The role of zinc in Alzheimer’s disease. International Journal of Alzheimer’s disease. 2011

28 Kaur, A., Kumar, S., Jaiswal, N., Vashisht, A., Kumar, D., Gahlay, G. K. and Mithu, V. S. (2019) NMR characterization of conformational fluctuations and noncovalent interactions of SUMO protein from Drosophila melanogaster (dSmt3). Proteins: Structure, Function, and Bioinformatics

29 Garg, A., Manidhar, D. M., Gokara, M., Malleda, C., Reddy, C. S. and Subramanyam, R. (2013) Elucidation of the binding mechanism of coumarin derivatives with human serum albumin. PLoS One. 8, e63805

30 Schägger, H. (2006) Tricine–sds-page. Nature protocols. 1, 16

31 Vranken, W. F., Boucher, W., Stevens, T. J., Fogh, R. H., Pajon, A., Llinas, M., Ulrich, E. L., Markley, J. L., Ionides, J. and Laue, E. D. (2005) The CCPN data model for NMR spectroscopy: development of a software pipeline. Proteins: Structure, Function, and Bioinformatics. 59, 687–696

32 Lin, D.-Y., Huang, Y.-S., Jeng, J.-C., Kuo, H.-Y., Chang, C.-C., Chao, T.-T., Ho, C.-C., Chen, Y.-C., Lin, T.-P. and Fang, H.-I. (2006) Role of SUMO-interacting motif in Daxx SUMO modification, subnuclear localization, and repression of sumoylated transcription factors. Molecular cell. 24, 341–354

33 Mielke, S. P. and Krishnan, V. V. (2009) Characterization of protein secondary structure from NMR chemical shifts. Progress in nuclear magnetic resonance spectroscopy. 54, 141

34 Baxter, N. J. and Williamson, M. P. (1997) Temperature dependence of 1 H chemical shifts in proteins. Journal of biomolecular NMR. 9, 359–369

35 Baxter, N. J., Hosszu, L. L., Waltho, J. P. and Williamson, M. P. (1998) Characterisation of low free-energy excited states of folded proteins. Journal of molecular biology. 284, 1625–1639

36 Williamson, M. P. (2003) Many residues in cytochrome c populate alternative states under equilibrium conditions. Proteins: Structure, Function, and Bioinformatics. 53, 731–739

37 Kumar, A., Srivastava, S. and Hosur, R. V. (2007) NMR characterization of the energy landscape of SUMO-1 in the native-state ensemble. Journal of molecular biology. 367, 1480–1493

38 Tunnicliffe, R. B., Waby, J. L., Williams, R. J. and Williamson, M. P. (2005) An experimental investigation of conformational fluctuations in proteins G and L. Structure. 13, 1677–1684

39 Cierpicki, T. and Otlewski, J. (2001) Amide proton temperature coefficients as hydrogen bond indicators in proteins. Journal of biomolecular NMR. 21, 249–261

40 Song, J., Durrin, L. K., Wilkinson, T. A., Krontiris, T. G. and Chen, Y. (2004) Identification of a SUMO-binding motif that recognizes SUMO-modified proteins. Proceedings of the National Academy of Sciences. 101, 14373–14378

41 Kaur, A., Kumar, S., Jaiswal, N., Vashisht, A., Kumar, D., Gahlay, G. K. and Mithu, V. S. (2019) NMR characterization of conformational fluctuations and noncovalent interactions of SUMO protein from Drosophila melanogaster (dSmt3). Proteins: Structure, Function, and Bioinformatics. 87, 658–667

42 Sapienza, P. J. and Lee, A. L. (2010) Using NMR to study fast dynamics in proteins: methods and applications. Current opinion in pharmacology. 10, 723–730

43 Hecker, C.-M., Rabiller, M., Haglund, K., Bayer, P. and Dikic, I. (2006) Specification of SUMO1- and SUMO2-interacting motifs. Journal of Biological Chemistry. 281, 16117–16127

